# Optimization of subsampling, decontamination, and DNA extraction of difficult peat and silt permafrost samples

**DOI:** 10.1101/2020.01.02.893438

**Authors:** Alireza Saidi-Mehrabad, Patrick Neuberger, Maria Cavaco, Duane Froese, Brian Lanoil

**Affiliations:** Department of Biological Sciences, University of Alberta, Edmonton, Alberta, T6G 2E9, Canada; Department of Earth and Atmospheric Sciences, Edmonton, Alberta, T6G 2E3, Canada

**Keywords:** Permafrost, Subsampling and decontamination, DNA extraction, 16S rRNA gene PCR, Contamination detection, Permafrost Microbiome

## Abstract

This study aims to act as a methodological guide for contamination monitoring, decontamination, and DNA extraction for peaty and silty permafrost samples with low biomass or difficult to extract DNA. We applied a biological tracer, either only in the field or both in the field and in the lab, via either spraying or painting. Spraying in the field followed by painting in the lab resulted in a uniform layer of the tracer on the core sections. A combination of bleaching, washing, and scraping resulted in complete removal of the tracer leaving sufficient material for DNA extraction, while other widely used decontamination methods did not remove all detectable tracer. In addition, of four widely used commercially available DNA extraction kits, only a modified ZymoBIOMICS™ DNA Microprep kit was able to acquire PCR amplifiable DNA. Permafrost chemical parameters, age, and soil texture did not have an effect on decontamination efficacy; however, the permafrost type did influence DNA extraction. Based on these findings, we developed recommendations for permafrost microbiologists to acquire contaminant-free DNA from permafrost with low biomass.

**IMPORTANCE:** Permafrost has the capacity to preserve microbial and non-microbial genomic material for millennia; however, major challenges are associated with permafrost samples, including decontamination of samples and acquiring pure DNA. Contamination of samples during coring and post coring handling and processing could affect downstream analyses and interpretations. Despite the use of multiple different decontamination and DNA extraction methods in studies of permafrost, the efficacy of these methods is not well known. We used a biological tracer to test the efficacy of previously published decontamination methods, as well as a bleach-based method we devised, on two chemically and structurally different permafrost core sections. Our method was the only one that removed all detectable tracer. In addition, we tested multiple DNA extraction kits and modified one that is able to acquire pure, PCR amplifiable DNA from silty, and to some extent from peaty, permafrost samples.

## INTRODUCTION

Permafrost, i.e. Earth materials below 0°C for at least two years and up to millions of years, acts as an archive of past environments and ecosystems, preserving biological material as a result of its isolation from atmospheric inputs, low temperatures, and low water activity (1). Ancient DNA derived from long-dead organisms is an important example of such material and has been used for a variety of purposes, ranging from reconstructing human migration patterns to reconstituting the genomes of extinct organisms such as the woolly mammoth and North American horses (2-6). Furthermore, permafrost-dwelling microbes may also play important roles in carbon cycling by conversion of permafrost organic carbon to methane and carbon dioxide, both important greenhouse gasses (7-10). The use of high-throughput sequencing technologies has enriched our understanding of microbial communities in permafrost and ancient DNA. However, these technologies require the extraction of high yields of DNA devoid of contaminants (11).

Obtaining DNA devoid of contaminants from environmental samples, especially from those with low biomass such as permafrost, is often challenging. Such samples are prone to external contamination during drilling and collection in the field and handling in the laboratory, which could lead to misinterpretation of microbial diversity, activity, or ancient DNA studies (12-14). External contamination is particularly problematic in DNA-based approaches due to the high sensitivity in detecting, amplifying and sequencing of DNA. Several methods have been used for permafrost decontamination, such as scraping the outer surface of cores, fracturing of cores followed by clean subsampling from the interior of the core sections (i.e. “disk sampling”), or washing the cores with DNase (e.g. (15-17); Table S1). Either scraping or disk sampling are the most commonly used protocols (Table S1); however, the efficacy of these methods in removing external contaminants is not well characterized (see, for example, (12)).

Ancient DNA (aDNA) and deep subsurface (both sediment and ice) microbiology studies face similar challenges to permafrost DNA studies, with high potential for contamination due to low endogenous cell and DNA abundance in the samples. Such studies have formalized highly stringent sampling and decontamination protocols, with protocols to minimize contamination and controls to monitor contamination at all stages from sampling to downstream analyses (e.g. (18-24)). Similar approaches may be beneficial for permafrost studies. For example, a unique tracer or combination of tracers added during drilling is used to monitor contamination in deep subsurface microbiology studies (25). Similar tracers have also been used in permafrost microbiology, but only rarely (e.g. (26-28)). Likewise, many decontamination methods have been systematically tested for ancient DNA studies of skeletal remains. Some of these methods, such as scraping (29) and disk sampling (30), have been used for permafrost decontamination as well. However, other methods used for aDNA studies of bone, including UV irradiation (31), and treatment with household bleach (32) have not been tested on permafrost intended for microbial work. Bleaching, in particular, has proven to be highly effective in removing external contaminants without damaging the genomic material within the samples in both ancient remains and ice cores (20, 32-34).

Another major issue in permafrost molecular studies is low DNA yield and poor quality of isolated DNA due to co-extracted chemical inhibitors (35-37). Permafrost researchers have utilized either commercial DNA extraction kits, most of which are based on mechanical disruption followed by DNA purification, or chemical DNA extraction protocols. Commercial mechanical disruption-based kits provide consistent DNA yield (although yield differs significantly between kits) and similar community composition, while chemical DNA extraction approaches are less consistent (38). Issues with co-extraction of chemical inhibitors have led some researchers to add extra purification steps. In some cases (e.g. 39-41), additional purification can lead to a loss of DNA or biases in the evaluation of community structure, although observable bias is not always seen (e.g. 38). To our knowledge, there have been no comparative studies assessing the efficacy of commercial kits for DNA extraction of difficult permafrost samples of different textures and chemistry.

In this study, we tested the efficacy of several decontamination methods on permafrost with the aid of a microbial tracer. In addition, we compared DNA yield and purity for four widely used commercially available soil DNA extraction kits with peaty and silty permafrost samples, with and without modifications of the manufacturer’s protocol. We developed recommendations for permafrost researchers for sample handling and processing, contamination detection and control, and DNA extraction.

## MATERIALS AND METHODS

### Site description and sampling strategies

A 3.97 m long, 10 cm diameter continuous permafrost core (termed DHL-16) was collected in May, 2016 adjacent to cores collected and presented previously (2 m lateral; GPS: 65.21061N, and 138.32208W; Fig. 1) (42). Two intervals of this core were sampled for this study. The first was a lower silt unit (from a depth of 254 cm to 336 cm, here called DH-2) dating to the Pleistocene, between 11,650 and 15,710 cal yr BP based on radiocarbon dating and age modelling (42). The second was an upper peat unit (from a depth of 105 cm to 212 cm, here called DH-1) dating to the early Holocene between 8,190 and 10,380 cal yr BP. The organic/silt boundary was determined at 244 cm (∼10,400 cal yr BP), placing the DH_2 core segment right around the start of the Holocene geological epoch.

**FIGURE 1.**
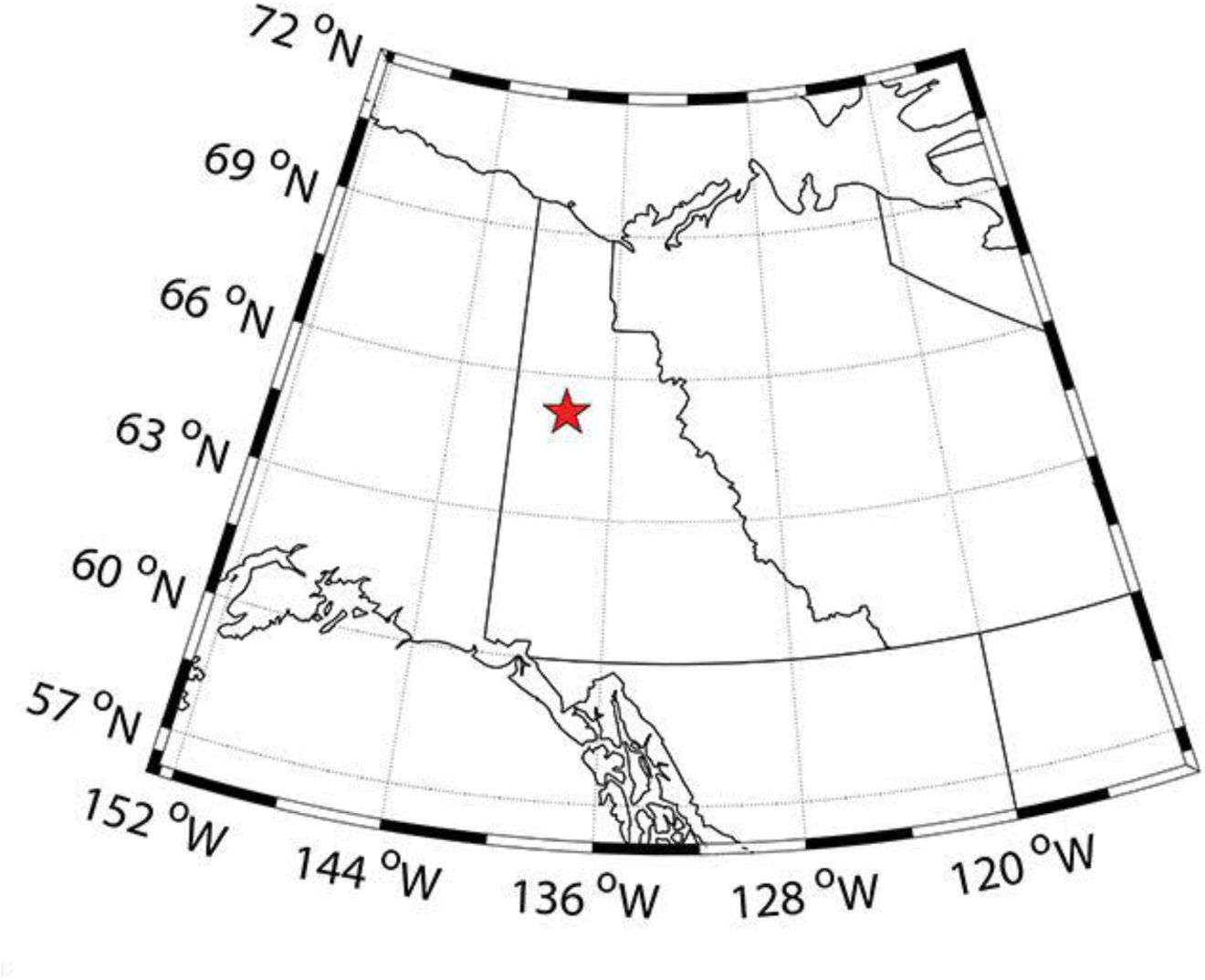
Map of Yukon region showing the coring location (DHP174) for DHL_16 core. The location was within the continuous permafrost zone (90-100% permafrost extent).

The surface material at our sampling site was approximately 2.5 m below the surface of surrounding undisturbed sites. To access the frozen permafrost table, we removed approximately 10-20 cm of thawed material with a shovel. The core was extracted by vertical drilling with a gas-powered drill with a custom-made diamond bit. Upon removing the core segments from the core catcher, the organic materials stuck to the surface of the core were scraped off with a clean pocket knife and the core was immediately sprayed with our contamination tracer (see below). Frozen core segments were placed in heat-sealed clear plastic bags (ULine, Canada), placed in coolers with ice packs for the duration of coring, and then stored at −20°C during transportation and subsequent analyses. At the University of Alberta, the DH_1 and DH_2 core segments were cut vertically into ⅓ and ⅔ subsections with the aid of a masonry saw. The ⅔ section was used to test decontamination and DNA extraction protocols, while the ⅓ section was used for chemical analyses

### Contamination tracer

Approximately 2.8 × 10^7^ cells/ml of *Escherichia coli* strain DH10B harboring a pBAD vector (Thermofisher Scientific, Canada), suspended in a total of 50 ml 1× PBS, was sprayed from a spray bottle on the core catcher, diamond bits, and the surface of the frozen cores (43-45). pBAD is an expression vector that codes for the mNeonGreen protein. This vector and its product was targeted as the main contamination tracer in this study via PCR of vector sequences and macro-photography of the mNeonGreen protein fluorescence under 470 nm wavelength using a xenon arc lamp (Sutter Instruments; California, USA).

### Sterilization procedure of the tools and work areas

To maintain cleanliness in the sub-sampling laboratory environment, we followed recommendations for ancient DNA and deep subsurface microbiological work (21, 46). These recommendations include the use of Tyvek clothing covers, masks, and gloves; sterilization of all equipment via baking, bleaching, or both; subsampling in a class 1000 clean space with no history of DNA extraction or PCR amplification of DNA; and monitoring of the space for potential contaminants. For full details, see supplemental methods.

### Basic chemical parameter analyses of the core segments

The ⅓ core sections of DH_1 and DH_2 were cut into 1-cm^3^ cubes with a handsaw in a 4°C cold room. The analyses of water content, organic carbon content, and pH were determined based on standard methods (see supplemental methods for details).

### Decontamination and subsampling methods

To prepare the samples for intentional contamination and decontamination, the ⅔ section of each core segment was cut horizontally into multiple disks (Fig. S2). Except for the piece selected for the decontamination protocol g (see below), one side of the disks was painted with a total of 5.3 × 10^8^ cells ml^-1^ of *E.coli* with pBAD suspended in 1× PBS using a 25 mm paintbrush. The other side was not painted in the laboratory and thus any spike present was the result of spraying in the field. For decontamination protocol g, the disk was cut into three rectangular subsections (Fig. S2). One rectangular piece was painted with the spike as above, another painted with a total of 18 µg of pBAD vector DNA isolated using QIAprep® Spin Miniprep Kit, by the manufacturer’s instructions (MO Bio, Qiagen Canada), and the third piece was not painted in the laboratory (Fig. S2).

Seven decontamination methods were tested in this study: a) scraping off external, potentially contaminated material by shaving the exterior of the cores 4-5 times with a series of 0.012”/0.30 mm single edge blades (i.e. “scraping”, modified from (17)). b) Sampling of a fresh, uncontaminated face with brass fitters (1/2” O.D. × 1.2” O.D.) (similar to (16) with the only difference being that brass fitters were used instead of a stationary drill press (i.e. “disk sampling”)); c) disk sampling as in protocol b, but using a soil press for volumetric subsampling (similar to (47)). Protocol c was performed with a set of high-pressure 30 cm long and 1.5 mm thick stainless steel tubing. d) Disk sampling as in protocol b, but using a hammer, chisel and a hand saw to remove the outer, contaminated material (similar to (48), with the only difference being that a manual hand saw was used instead of an electric jigsaw, no clamps were utilized, and the cores were not cut into cubes). e) A combination of scraping and disk sampling with chisels and blades (similar to (35), with the only difference being that single edge blades were used instead of knives). f) UV irradiation of the disk (modified from (29)). In protocol f, a disk was placed in a clean, closed UV box (UVP C-70G Chromato-Vue® Cabinet; Analytik Jena, USA) ∼6 cm from the UV lamp of 15 watts and was subjected to UV light at 254 nm for 5 min (8.45 ×10^−17^ J/m^2^ UV dosage), 10 min (1.69 ×10^−16^ J/m^2^ UV dosage), 20 min (3.38 ×10^−16^ J/m^2^ UV dosage) or 30 min (5.07 ×10^−16^ J/m^2^ UV dosage) intervals. g) Scraping and bleaching (developed for this study based on (32, 33)). In protocol g, the surface of the rectangular piece was first washed with pre-chilled (4°C), full strength concentrated household bleach solution with no phosphorus compounds. Bleach was rinsed off with pre-chilled (4°C) Milli-Q water. The resulting loosened surface materials were removed via scraping with 0.012”/0.30 mm heavy duty single edge blades (Richard Ltd, Quebec Canada). This entire process was then repeated a second time. Unlike protocols a-f, protocol g started with subsampling first and then decontamination (Fig. S2).

Decontaminated samples were stored at −20°C prior to DNA extraction. Decontaminated samples obtained from protocols (a and c-f) were crushed into smaller pieces with a sterile chisel and hammer and prior to DNA extraction, they were allowed to thaw at room temperature. Thawed material was homogenized by mixing and the resulting material was subsampled for DNA work. Soils in brass fitters obtained via protocol (b) were left at room temperature prior to DNA extraction to allow easy removal of the material with the aid of a sterilized spatula and were later mixed and subsampled.

### DNA extraction

We compared seven DNA extraction protocols: four commercially available, well-established soil DNA extraction kits as recommended by the manufacturers as well as modifications to three of these commercial kits (see supplemental material for details of the modifications). The protocols used for soil DNA extraction were as follows: 1) Fast DNA™ SPIN kit for soil (MP Biomedicals, California, USA) by the manufacturer’s protocol; 2) Fast DNA™ SPIN kit for soil (MP Biomedicals, California, USA) with modifications; 3) OMEGA E.Z.N.A soil DNA kit (OMEGA-Bio-Tek, Georgia, USA) by the manufacturer’s protocol; 4) Powersoil^®^ Isolation kit (MO Bio Laboratories/Qiagen, Canada) by the manufacturer’s protocol; 5) Powersoil^®^ Isolation kit (MO Bio Laboratories/Qiagen, Canada) with modifications; 6) ZymoBIOMICS™ DNA Microprep kit (Zymo Research, California, USA), by the manufacturer’s protocol; and 7) ZymoBIOMICS™ DNA Microprep kit (Zymo Research, California, USA) with modifications. DNA yield was determined using a Qubit fluorometer device (Invitrogen. Ontario, Canada) via Quant-iT dsDNA HS Assay Kit (Invitrogen, Canada), calibrated using the manufacturer’s protocol. DNA was extracted from triplicate 1 g subsamples for DH_1 and DH_2, and triplicate 0.5 g subsamples from the control soil (CS; see below). A positive control soil sample (termed CS in this manuscript) was used to test the efficiency of each DNA extraction protocol (mentioned below) in obtaining contaminant free and PCR amplifiable DNA from a non-permafrost sample. The CS sample was an 8:1 ratio of peat and mineral subsoils from the rhizosphere of *Populus tremuloides*, mixed using a clean cement mixer for 10 minutes. Two blank negative controls with no soil added were prepared from each kit to trace possible contamination originating from kit reagents.

### Contamination detection

To determine if a decontamination procedure was successful, the isolated DNA was tested for the presence of pBAD-vector via PCR (see supplemental methods for PCR protocol details).

### 16S rRNA gene-targeted PCR protocol

16S rRNA genes were PCR amplified from the DNA obtained from the decontaminated samples to test for their PCR amplifiability, here used as a proxy for DNA purity. For more information, refer to supplemental methods.

## RESULTS

### Chemical characteristics of core sections DH_1 and DH_2

The DH_1 segment was a peaty unit with high organic matter content (mean = 95.7% w/w dried (±1.82%), n = 23), high gravimetric water content (mean = 91.8% w/w (±3.01), n = 23), and low pH (mean = 3.68 (±0.102), n = 24). DH_2 segment was a silty unit with lower organic matter content (mean = 39.58% w/w dried (±21.85%), n = 21), lower gravimetric water content (mean = 74.48% w/w (±18.02%), n = 21), and higher pH (mean = 6.04 (±0.54), n = 30) (Figure S1). The organic matter content range for DH_1 was relatively consistent (90.56% - 98.02%) (Fig S1). However, DH_2 varied widely in organic content (9.91% - 68.36%). A similar trend was observed regarding gravimetric water content, with the DH_1 fairly consistent (80.88-98.33%), but DH_2 samples varying dramatically (8.15-97.61%). pH did not change significantly in DH_1 with depth; however, the pH increased significantly with depth for DH_2, from 5.17 to 6.9 (Fig. S1).

### Decontamination testing

To test our decontamination protocol, we applied *E. coli* carrying a mNeonGreen protein expression vector to our core sections as a tracer. The tracer was applied by spraying the corer and the core sections in the field and/or by painting the core sections in the lab. Painting of the tracer on the core sections showed a uniform distribution of cells based on fluorescence of mNeonGreen protein as well as consistent amplification of the pBAD vector PCR product from all samples prior to decontamination (data not shown). The side of the disk where tracer was only applied in the field resulted in patches of spike and inconsistent amplification of the vector. However, the crystallized ice from the interior of the bags used for transporting the core sections always showed positive PCR amplification of the vector, indicating that the tracer was easily removed from the surface of the core. In addition, we noticed cutting the samples and handing in the lab resulted in the loss of the contamination tracer. Hence, we recommend the application of the tracer by painting prior to decontamination, as well as field application by spraying, to ensure decontamination is as thorough as possible.

Of the seven decontamination methods, scraping (protocol a) and UV irradiation (protocol f) retained the most material for subsequent biological work (Table 1). Conversely, disk decontamination with brass fitters (protocol b) was the most destructive, resulting in a very small quantity of decontaminated material. The soil press method (protocol c) did not perform well, resulting in crushing and thawing of the disk and bending of the tubes. Protocols d (disk sampling with chisel removal of outer material), e (disk sampling with scraping), and g (scraping and bleaching) resulted in a moderate quantity of samples for biological work (Table 1).

**TABLE 1.** Decontamination methods on permafrost samples DH_1 and DH_2.

| Protocol<sup>a</sup> | Fraction retained (mass %) |  | Colonies (<1 m away) |  | Colonies (> 1 m away)<sup>e</sup> |  | pBAD amplification<sup>f</sup> |  |
| --- | --- | --- | --- | --- | --- | --- | --- | --- |
|  | DH_1 | DH_2 | DH_1 | DH_2 | DH_1 | DH_2 | DH_1 | DH_2 |
| a | 94 | 92 | +<sup>d</sup> | + | +/- | +/- | + | + |
| b | 7 | 6 | + | + | +/- | +/- | - | - |
| c<sup>b</sup> | 0 | 0 | n.d.<sup>d</sup> | n.d. | +/- | +/- | +/- | +/- |
| d | 40 | 47 | + | + | +/- | +/- | +/- | +/- |
| e | 37 | 45 | + | + | +/- | +/- | +/- | +/- |
| f | 100 | 100 | + | + | +/- | +/- | +/- | +/- |
| g<sup>c</sup> | 40 | 32 | -<sup>f</sup> | - | - | - | - | - |

<sup>a</sup> Protocols: a = scraping, b = disk sampling with brass fitters, c = disk sampling with a soil press, d = disk sampling with a chisel, hammer and a hand saw, e = combination of scraping and disk sampling, f = UV irradiation, g = bleach and scraping. See methods for details.
<sup>b</sup> This protocol failed to acquire any samples due to bending of the tubing.
<sup>c</sup> NOTE: In this protocol, permafrost is first subsampled and then decontaminated; for other protocols, decontamination occurs before subsampling (see figure S2).
<sup>d</sup> n.d. = not done, + = detected, - = not detected, +/- = inconsistent detection.
<sup>e</sup> Colonies formed on nutrient rich media plates placed near work station.
<sup>f</sup> PCR amplification of the pBAD plasmid carried by the intentional contaminant. Indicates contamination.

The DNA from the soil samples was extracted via DNA extraction protocol 7 and tested via PCR of the pBAD vector. The decontaminated samples from protocols b and g were devoid of PCR amplifiable pBAD vector, indicating effective decontamination (Table 1). Decontaminated samples from protocols (a) and (c-f) resulted in amplification of the pBAD vector when tested with PCR, indicating incomplete decontamination. Protocols a-f resulted in colony formation on growth media left open in the room during decontamination, indicating contamination of the local environment. Such contamination could lead to subsequent cross-contamination of other samples. Protocol (g) was the only method that did not show colony formation on nearby growth media (Table 1). Thus, protocol (g) provided complete decontamination and a moderate amount of decontaminated material remaining, and therefore seems to be the best decontamination protocol for permafrost samples and was used for subsequent DNA extraction testing.

### DNA extraction testing

Following decontamination with protocol (g), DNA was extracted from the two permafrost samples as well as a positive control temperate soil (CS). The kits and protocols tested displayed varying efficiency and effectiveness in extracting DNA (Table 2). Protocol 1 did not result in any detectable DNA when it was used on either permafrost sample, but it resulted in the highest yield of DNA from CS (Table 2). Detectable, but low, DNA yield from DH_2 was obtained with Protocol 2 and Protocol 5; however, neither of these protocols provided detectable DNA from DH_1 (Table 2). Protocol 3 resulted in DNA yield from DH_1, DH_2, and CS (Table 2). Protocol 6, in contrast to other methods, was able to obtain detectable DNA from DH_1 and CS, but not DH_2 (Table 2). Protocol 7 produced DNA from both DH_1 and DH_2 (Table 2). All of the DNA extraction protocols provided high yields of DNA for the positive control temperate soil, (CS). The CS samples provided 2-3 orders of magnitude more DNA (47× – 754×) than the permafrost samples, no matter which extraction protocol was utilized (Table 2). Protocol 3 consistently resulted in PCR amplification from blank extractions, both with different kit lot numbers and different researchers; as a result, we did not test this protocol further (Table 2). We tested the purity of DNA obtained from unmodified kit protocols (i.e. protocols 1, 4, and 6) on the CS soil; all kits provided DNA pure enough to PCR amplify 16S rRNA genes. However, on permafrost soils, DNA from protocols 1 and 2 was not PCR amplifiable for either permafrost sample (Table 2). Several protocols gave differential results for the two different samples, with protocols 4 and 5 showing better PCR amplifiability with DH_2 and protocol 6 showing better PCR amplifiabilty with DH_1 (Table 2). For protocols 4 and 6, DNA yield was below the detection limit; however, PCR product was obtained (Table 2). Only protocol 7 provided consistently strong PCR amplification from both permafrost samples (Table 2).

**TABLE 2.** DNA extraction protocol on samples DH_1, DH_2, and CS.

| Protocol<sup>a</sup> | DNA yield (ng/g)<sup>b</sup> |  |  |  | PCR amplification |  |  |  |
| --- | --- | --- | --- | --- | --- | --- | --- | --- |
|  | DH_1<br>(±SD) | DH_2<br>(±SD) | CS<br>(±SD) | Kit<br>blank<sup>c</sup><br>(±SD) | DH_1 | DH_2 | CS | PCR<br>blank |
| 1 | BDL | BDL | 6633<br>(2310) | BDL | -<sup>d</sup> | - | +++ | - |
| 2 | BDL | 2.5<br>(0.1) | n.d. | BDL | - | - | n.d.<sup>d</sup> | - |
| 3 | 24.6<br>(0.2) | 31.8<br>(0.2) | 1517<br>(16) | 60.1<br>(3) | +<sup>d</sup> | + | + | + |
| 4 | BDL | 10<br>(0.1) | 660<br>(6.4) | BDL | + | ++<sup>d</sup> | ++ | - |
| 5 | BDL | 4.6<br>(0.4) | n.d. | BDL | - | ++ | n.d. | - |
| 6 | 0.7<br>(0.1) | BDL | 513<br>(14) | BDL | +++<sup>d</sup> | + | + | - |
| 7 | 1.1<br>(0.2) | 17<br>(0.2) | n.d. | BDL | +++ | +++ | n.d. | - |

<sup>a</sup>DNA extraction protocols: 1 = Fast DNA SPIN kit for soil (manufacturer's protocol), 2 = Fast DNATM SPIN kit for soil (modified), 3 = OMEGA E.Z.N.A soil DNA kit (manufacturer's protocol), 4 = Powersoil Isolation kit (manufacturer's protocol), 5 = Powersoil Isolation kit (modified), 6 = ZymoBIOMICSTM DNA Microprep kit (manufacturer's protocol), and 7 = ZymoBIOMICSTM DNA Microprep kit (modified).
<sup>b</sup>DNA was extracted from triplicate 1 g subsamples for DH\_1 and DH\_2, and triplicate 0.5 g subsamples from CS.
<sup>c</sup>Measured in ng of DNA.
<sup>d</sup>BDL = below the detection limit, n.d. = not done, + = weak PCR band, ++ = medium PCR band, +++ = strong PCR band, - = not detected.

## DISCUSSION

Deep subsurface microbiology studies have demonstrated the importance of contamination detection through the use of tracers (18, 49). Fluorescent latex beads similar in size to microbes have been used extensively in deep subsurface microbiology (e.g. (50)) and to a lesser extent in permafrost studies (e.g. (12, 51)) to track potential contamination during sample acquisition. However, these beads do not mimic microbes well (12, 52), are subject to quenching and bleaching of fluorescence (22), are labor-intensive to detect (53), and cannot be detected easily at low levels of contamination (19). Biological tracers have two major advantages relative to beads: they are biological particles and thus mimic contaminants better and they can be easily detected at very low levels by PCR (51). Intact cells that are not found in permafrost that carry a well-characterized target DNA molecule, such as a plasmid, are an ideal contamination tracer. In this study, we utilized *E.coli* mNeonGreen-expressing cells, which are a commercial product and thus are not found in permafrost. This tracer can be visualized by fluorescence of the mNeonGreen protein and the pBAD plasmid is easily detected at low levels by PCR.

Applying the tracer to the wrong sampling component or at the wrong time may lead to a false negative, i.e. the presumption that decontamination is complete when the lack of detection of the tracer is actually due to loss during handling (12, 18, 51). Based on our observations, tracer should be applied both to the drilling apparatus and cores in the field and again in the laboratory; application solely in the field led to inconsistent detection of tracer even before decontamination. Furthermore, we found that applying the tracer by painting rather than by spraying provided a more consistent coverage of samples.

Our results showed that none of the tested decontamination methods were able to completely remove the tracer except the bleach wash method and disk sampling method with brass fitters. Bleach is cheap and readily available in comparison to costlier DNAse and RNAse decontamination solutions used in the past (15, 32). Bleach was effective in removing our tracer and left a moderate amount of the material available for subsequent work. In contrast, while the fitter based protocol used a clean subsampling approach, it yielded a low quantity of subsamples.

One possible disadvantage of using bleach for decontaminating permafrost segments is changes to the chemistry of the samples. We tackled this potential issue by splitting the core section into separate samples for chemistry and biology (i.e. ⅓ and ⅔ sections), which allowed preservation of samples for chemistry work, and a sufficient amount of material for decontamination and DNA extraction. However, if the amount of material available is restricted, this approach may not be tenable.

The rest of the tested methods, based on the most commonly used method in published permafrost studies (i.e. disk sampling or scraping; Table S1) resulted in inconsistent PCR amplification of the tracer from decontaminated samples. One possible reason for a lack of decontamination was physical contact of the clean interior pieces with contaminated materials and dust generation during the subsampling. We noted that our test plates were contaminated with tracers and other cells during disk sampling methods, likely indicating the production of contaminated dust or aerosols during processing, similar to previous findings (23, 54). Thus, methods that minimize dust and aerosol generation are recommended to decrease the possibility of re-contaminating cleaned samples.

In the case of scraping, insufficient removal of the contaminated surface of the core section may have been another reason for detecting the tracer. Bang-Andreasen and colleagues (12) demonstrated that their intentional contamination spike was still detectable down to 17 mm depth after coring; thus, scraping, which in our experiment only removed 2-3 mm after 4-5 scrapes, is insufficient to decontaminate the core. The ineffectiveness of the scraping method has also been reported in ancient DNA studies (32). Thus, we strongly recommend against scraping as the sole decontamination method for permafrost cores.

In our experiment, commercial DNA extraction kits vary in both DNA yield and purity. In a previous study, the Fast DNA™ SPIN kit for soil (MP Biomedicals, California, USA) provided the highest DNA yield from permafrost, although it required further purification (38). However, in our experiment while the Fast DNA™ SPIN kit for soil gave the highest yield in the control soil, no detectable DNA was obtained from the permafrost. The modified protocol for ZymoBIOMICS™ DNA Microprep kit (Zymo Research, California, USA) was the only protocol able to yield sufficient PCR amplifiable DNA. It is unclear whether the same kit or the same modifications will always provide optimal results; thus, when there is sufficient sample, we recommend testing of several commercially available kits and modification of those protocols (e.g. see supplemental methods) to obtain the maximum amount of pure DNA from permafrost.

It is critical to utilize DNA extraction blank controls since the kit reagents could introduce contamination. In one protocol, the negative control for the kit always showed amplification, indicating contamination from the kit reagents. Contamination via kit reagents has been observed in other studies as well (e.g. (55)). Eisenhofer and colleagues have noted and summarized some species from a large variety of microbiome studies that are regularly found in DNA extraction kits. Thus, it is clear that extraction kits can and often do introduce contaminants: kits should be selected with care for low biomass samples such as permafrost that are prone to contamination. Furthermore, extractions should include extensive positive (control soils) and negative (blank extraction) controls.

Our results indicate that basic soil chemical parameters did not influence the spike penetration or decontamination procedures; however, these parameters did affect DNA extraction yield. The silty core generally provided a higher DNA yield than the peaty core, indicating that permafrost chemical and physical parameters can affect DNA extraction.

## CONCLUSIONS AND RECOMMENDATIONS

We recommend the following to prevent contamination of permafrost samples intended for microbial work:

1. A biological spike should be applied both in the field via spraying and in the lab by painting of the core sections. The spike should be allowed to fully freeze onto the core. PCR should be used to detect the applied biological tracer: clean samples should be negative; removed material should be positive.
2. Ancient DNA protocols for sample handling should be followed whenever possible (e.g. (21, 46)). These protocols were developed to minimize external contamination and cross-contamination between samples. These protocols are evolving and should be updated regularly. We have provided a summary of these guidelines in this manuscript (see supplemental methods for details)
3. Combined bleach wash and shaving are the most effective method for decontaminating permafrost samples intended for DNA work. We recommend against utilizing only disk decontamination or scraping, as these approaches did not remove our tracer.
4. Multiple DNA extraction kits should be tested for the specific samples, with both positive (temperate soil) and negative (reagents only) controls. In our experiment, modified ZymoBIOMICS™ DNA Microprep kit (Zymo Research, California, USA) was the most effective method in extracting DNA from silty permafrost and to some extent from peaty permafrost; however, other samples may respond better to other DNA extraction protocols.

## ACKNOWLEDGMENTS

We would like to thank Dr. Lauren Davies for assistance with radiocarbon dating and chronology of the cores, Ali Naeimi Nezamabad for assistance with the sampling location map, Sasiri Bandara, Casey Buchanan, and Joseph Young for assisting us in obtaining the permafrost core segments, and Tania Strilets for editing the current manuscript. We would like to extend our thanks to the University of Alberta (UANRA grant) and Polar Knowledge Canada (NSTP grant) for their student financial support. We would like to thank the following people for their insightful comments, support and allowing us to use their equipment: Dr. Martin Sharp, John Sherman, Suzan Gater, Richard Mah, and Dr. Alberto Reyes. We are grateful to Zymo Research and Cedarlane^®□^ labs for their generous donation of the DNA extraction kits and guidance.

